# Past climate changes, population dynamics and the origin of Bison in Europe

**DOI:** 10.1101/063032

**Authors:** Diyendo Massilani, Silvia Guimaraes, Jean-Philip Brugal, E. Andrew Bennett, Malgorzata Tokarska, Rose-Marie Arbogast, Gennady Baryshnikov, Gennady Boeskorov, Jean-Christophe Castel, Sergey Davydov, Stephane Madelaine, Olivier Putelat, Natalia Spasskaya, Hans-Peter Uerpmann, Thierry Grange, Eva-Maria Geigl

## Abstract

Climatic and environmental fluctuations as well as anthropogenic pressure have led to the extinction of much of Europe’s megafauna. Here we show that the emblematic European bison has experienced several waves of population expansion, contraction and extinction during the last 50,000 years in Europe, culminating in a major reduction of genetic diversity during the Holocene. Fifty-seven complete and partial ancient mitogenomes from throughout Europe, the Caucausus and Siberia reveal that three populations of wisent (*Bison bonasus*) and steppe bison (*B. priscus*) alternated in Western Europe correlating with climate-induced environmental changes. The Late Pleistocene European steppe bison originated from northern Eurasia whereas the modern wisent population emerged from a refuge in the southern Caucasus after the last glacial maximum. A population overlap in a transition period is reflected in ca. 36,000 year-old paintings in the French Chauvet cave. Bayesian analyses of these complete ancient mitogenomes yielded new dates of the various branching events during the evolution of Bison and its radiation with Bos that lead us to propose that the genetic affiliation between the wisent and cattle mitogenomes result from incomplete lineage sorting rather than post-speciation gene flow.

**Significance:** Climatic fluctuations during the Pleistocene had a major impact on the environment and led to multiple megafaunal extinctions. Through ancient DNA analyses we decipher these processes for one of the largest megafauna of Eurasia, the bison. We show that Western Europe was successively populated during the Late Pleistocene by three different bison clades or species originating from the Caucasus and North-Eastern Europe that can be correlated to major climatic fluctuations and environmental changes. Aurignacian cave artists were witnesses to the first replacement of bison species ~35,000 years ago. All of these populations went extinct except for one that survived into the Holocene where it experienced severe reductions of its genetic diversity due to anthropogenic pressure.

## Introduction

Drastic climatic fluctuations during the Pleistocene in the northern hemisphere led to population contractions, extinctions, reexpansions and colonizations of fauna and flora (1). Bison, along with other large ungulates thrived during the Middle and Late Pleistocene (2). Numerous cave paintings and engravings in France and Spain, such as those in the caves of Chauvet, Lascaux and Altamira, attest to the important role this impressive animal played for the late Paleolithic hunter-gatherers. The steppe bison (*Bison priscus* (Bojanus, 1827)) appears in the fossil record during the Early Middle Pleistocene, replacing another archaic but smaller forest-adapted bison (B. *schoetensacki(3)*), which went extinct ca. 700 kiloyears ago (kya) (4). Since *B. priscus* was adapted to the cold tundra-steppe, occurrence of its remains is considered indicative of open environments (5). It roamed over Europe and Asia, and also crossed the Bering Strait during the Middle Pleistocene to populate North America where it evolved into the American Bison *B. bison* (3, 6, 7). The numerous fossil remains display a pronounced sexual dimorphism, and a large initial body size, gradually decreasing throughout the Pleistocene (5). Differences in morphology related to climatic, environmental and topographic conditions have led several authors to propose a high diversity for the Pleistocene cold-steppe bison expressed as subspecies or ecomorphotypes (5, 6).

The taxonomy, evolutionary history and paleobiogeography of the genus *Bison* in Eurasia, and of the European bison or wisent *B. bonasus* (Linnaeus, 1758) in particular, is still patchy, despite a rich fossil record and its current endangered status (e.g., (8–11)). Indeed, two opposing hypotheses on the evolution of bison in Eurasia coexist (2). Traditionally, it has been considered that bison developed within one single phylogenetic line (B. *schoetensacki-B. priscus-B. bonasus*), but it has also been proposed that at least two parallel lines of bison existed, one being the line of forest bison from *B. schoetensacki* (Freudenberg, 1910) to the recent *B. bonasus* and the other being the line of the steppe bison *B. priscus* (for a review see (2)). Thus, the phyletic relationships between *B. shoetensacki, B. priscus* and *B. bonasus*, as well as the approximate date and geographical origin of the wisent remain elusive, due in part to the limited power of paleontological studies to resolve species-level mammalian taxonomy issues or detect broad-scale genetic transitions at the population level (12).

*B. priscus* disappeared from the fossil record of Western Europe at the end of the Pleistocene, around 12-10 kya, and relict populations of *B. priscus* seem to have survived until the beginning of the Middle Holocene (7-6 kya) in Siberia (i.e., (13, 14)). In Europe, *B. priscus* is believed to have been replaced at the end of the Pleistocene or during the Holocene by the morphologically (eidonomically) distinguishable wisent, *B. bonasus* (2, 10, 15, 16). At least two sub-species are recognized: (i) *B. b. bonasus* Linnaeus, 1758 from the Lithuanian lowland and the Polish Biatowieza ecosystem, and (ii) the Caucasian highland *B. b. caucasicus* (Turkin and Satunin, 1904). (17). *B. priscus* was adapted to forest-steppe and steppe, and *B. bonasus* to forest and mountain-forest environments. *B. priscus* and *B. bonasus* are anatomically much closer to each other than to other more ancient bison, such as *B. schoetensacki. B. bonasus* has a relatively more massive rear quarter and shorter horns compared to *B. priscus* which has longer and slightly curved horns and a smooth double-humped appearance (15, 16). *B. priscus* and *B. bison* (Linnaeus, 1758), which are grazers, have a lower head position than *B. bonasus*, which is a mixed feeder (18). It is, however, very difficult to assign fossil bison bones to either species (2). Cave paintings of bison from caves in France and Spain are often classified as belonging to either *B. priscus* or to *B. bonasus* (19). The diversity of the cave art depictions and the large range of their occurrence is interpreted as indicating an origin of *B. bonasus* in the area between southern Europe and the Middle East and of its existence well before the end of the Late Pleistocene at a time when *B. priscus* was still present (19).

Both extant bison species narrowly escaped extinction. The American *B. bison* was almost wiped out during the 19^th^ century through commercial hunting and slaughter, but also introduced bovine diseases and competition with domestic livestock (20). The wisent as well almost went extinct at the beginning of 20^th^ century. Indeed, similar to other large herbivores, such as the aurochs, intensification of agriculture since the Neolithic pushed the wisent into the forests of Eastern Europe (18) where it was strictly protected for a few centuries as a royal game animal (11). During the First World War, however, diminished population size followed by poaching led to its extinction in the wild (11). Of the 54 remaining wisents living in captivity at the beginning of the 1920s, the descendants of just 12 animals constitute the entire extant population (11).

The wisent is still poorly characterized genetically. While genetic markers from the autosomes and Y chromosomes of American bison and wisent are closer to each other than to the other members of the genus *Bos* and they can reproduce and give rise to fertile offspring, their mitochondrial genomes are phylogenetically separated (9, 21, 22). Indeed, mitochondrial sequences of the American bison and the yak *Poephagus mutus* f. *grunniens* (Linnaeus, 1758) form a distinct cluster while the wisent occupies a phylogenetic position closer to *Bos primigenius* f. *Taurus* (Linnaeus, 1758), a phenomenon that has been explained by incomplete lineage sorting or ancient hybridization (21, 22). European, Siberian and American *B. priscus* mitogenomes were shown to be phylogenetically closer to *B. bison* than to *B. bonasus* (7, 14, 23).

Ancient DNA (aDNA) studies have the potential to better resolve taxonomy than paleontological studies, in particular at the species level, and have revealed a far more dynamic picture of megafaunal communities, biogeography and ecology, including repeated localized extinctions, migrations, and replacements (e.g., (12) and citations therein). To better understand the phylogeography and evolution of the wisent and to reconstruct its origin, we performed a study of the mitochondrial genome over the past 50,000 years (50 kyrs) across Europe and Asia. Using Pleistocene skeletal remains from Siberia (Yakutia), the Caucasus and Western Europe (France, Switzerland) as well as Neolithic and Medieval samples from France, and also pre-bottleneck *B. bonasus* samples from the early 20^th^ century from the Polish lowland and the Caucasian highland lines, we constructed a phylogenetic framework based on both the hypervariable region of the mitochondrial DNA as well as complete mitochondrial genomes of selected specimens. Our results give new insights into the climate-driven dynamics of the bison in Europe.

## Results

### The evolution of the mitochondrial hypervariable region

We analyzed 57 paleontological and 25 historical specimens (see SI Table 1) targeting the mitochondrial hypervariable regions with four fragments of at most 150 bp. We obtained a 367 bp-long sequence from 43 specimens, and smaller sequences from 13 additional ones (SI Fig. 1). A Maximum Likelihood phylogenetic tree was constructed using these sequences (Fig. 1). Three clades can be clearly distinguished. The first clade *Bp*, which is more divergent from the other two, comprises samples from Pleistocene Yakutia (Siberia) and from southern France (Berbie and lower stratigraphic layers of Gral), dating from 44 to 26 kya and from 39 to 15 kya, respectively. Clade *Bp* corresponds to the *B. priscus* lineage previously described for Siberia, North America and Europe (7, 23). The French and Siberian sequences of this clade are phylogenetically close and lack a phylogeographic structure. This reveals that a relatively homogeneous population of steppe bison was distributed during the Late Pleistocene not only in Siberia and northern America but also throughout the entire northern part of the Eurasian continent up to its most-western part, France.

**Fig. 1.**
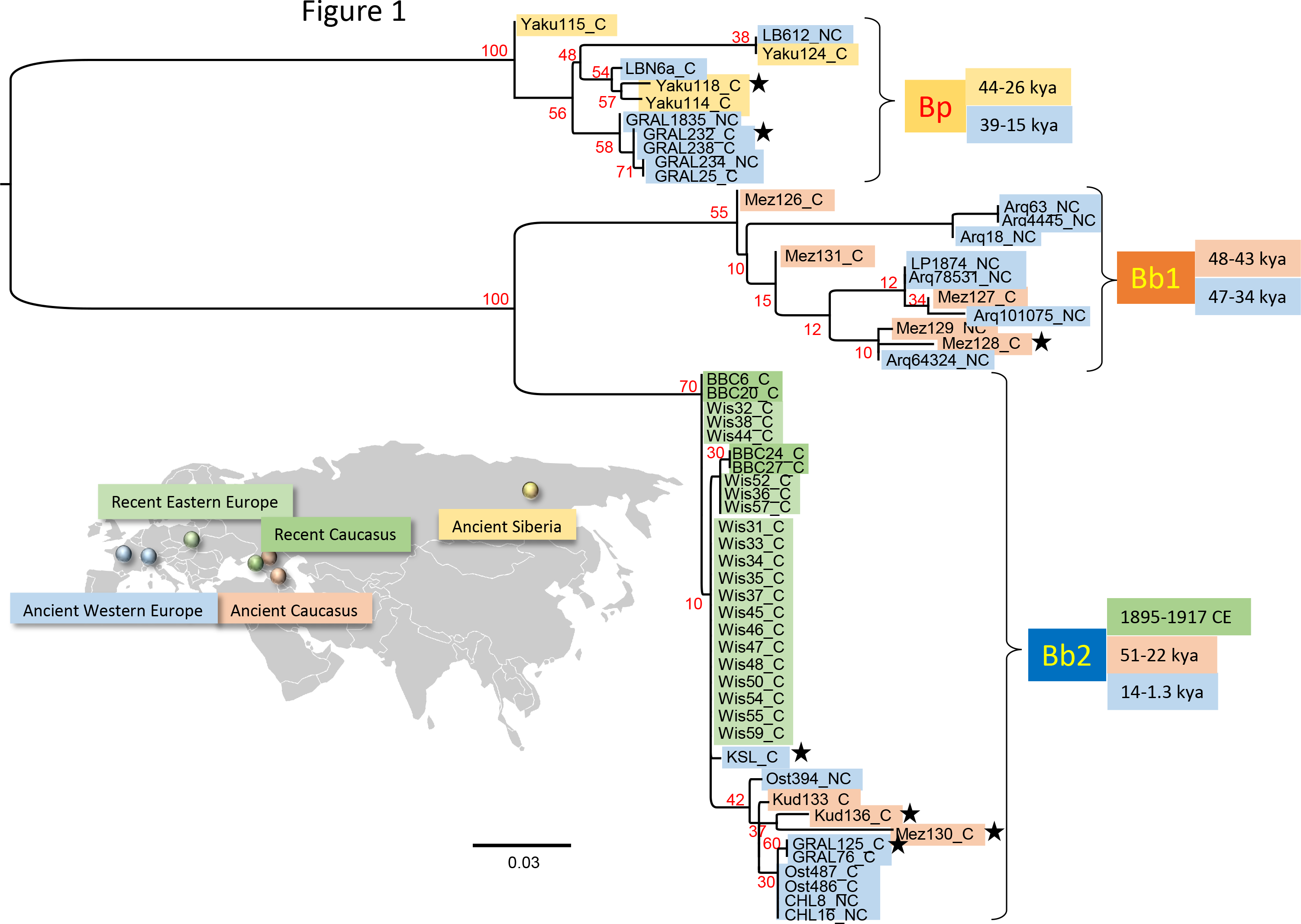
Maximum Likelihood phylogeny of the ancient Bison hypervariable region. ML analyses of the HVR produced in this study using PHYML, a HKY+I+G substitution model and 500 bootstraps. The bootstrap support of the nodes is indicated in red. The geolocalization of the analyzed samples is represented using a color code to distinguish five origins and time periods as represented on the Eurasiatic continent map (see SI. Fig.4 for a map of the distribution of the Western European sites). Three clades can be clearly distinguished, the *Bison priscus (Bp*) clade and two *Bison bonasus* clades (Bb1 and Bb2). The scale corresponds to the number of nucleotide substitutions per site. The samples that have allowed amplification of all 4 PCR fragments as represented in SI fig. 1 are indicated by the suffix _C whereas those for which one or more fragments were missing are indicated by the suffix _NC. The stars indicate the samples that were used for the full mitogenome analysis presented in Fig. 3.

The two other clades, *Bb1* and *Bb2*, are more closely related to the extant wisent *B. bonasus.* The *Bb1* clade, significantly divergent from *Bb2*, contains most of the specimens from the Mezmaiskaya cave Northwest of the Caucasus in Russia, dating from ca. 48 to 43 kya, as well as seven specimens from southern France (Arquet, Plumettes), dating from ca. 47 to 34 kya. Thus, the geographical range of the *Bb1* clade stretched from the northern Caucasus to the south of France at its latest, from 48 to 34 kya during the overall mild Marine Isotope Stage (MIS) 3 (ranging from 57 to 29 kya (24), http://www.lorraine-lisiecki.com/LR04MISboundaries.txt). None of the haplotypes of the *Bb1* clade was found in the more recent specimens, suggesting that the corresponding population may have been the first to disappear. This *Bb1* population in France was apparently replaced by the steppe bison of the *Bp* clade around 30 kya, the latter remaining there throughout the glacial period of MIS2 (ranging from 29 to 14 kya (24), http://www.lorraine-lisiecki.com/LR04MISboundaries.txt).

Clade *Bb2* includes both ancient specimens from the Caucasus and Western Europe as well as all recent and extant *B. bonasus* specimens. Nearly all ancient samples belonging to this clade are distinct from the more recent populations and include a ca. 49 ky-old specimen from the Mezmaiskaya cave, two specimens from the Kudaro cave in the central part of the southern slope of the Greater Caucasus dated at 38 and 22 kya, and specimens from Western Europe: one specimen from the Kesslerloch cave (Switzerland) dated at 14 kya, and, in France, two bison from the upper stratigraphic sequence of the Gral dated at 12 kya, two 5.2kyr-old bison samples from the Neolithic site of Chalain, as well as three medieval (7^th^-8^th^ century CE) specimens from Alsace. The members of this clade represent the western European Pleistocene-Holocene lineage of *B. bonasus* and display a high mitochondrial diversity. This lineage appears to have replaced the *B. priscus* lineage, at least in France, at the end of the Upper Pleistocene between 15 and 12 kya, coinciding with the onset of a more temperate climate, and to have persisted in France up to the Middle Ages. Apart from the sequence found in the sample from Kesslerloch (12.2 kya), none of the ancient Bb2 sequences are present in the extant mitochondrial gene pool.

Within the *Bb2* clade, a compact group of closely related sequences, comprise the Upper Pleistocene Kesslerloch specimen and the 1898-1917 pre-bottleneck wisents from both Poland and the Caucasus that almost became extinct at the end of the First World War. Five out of 24 of these pre-bottleneck bison have a hypervariable region (HVR) sequence identical to that of extant *B. bonasus*, whereas the rest differ from the extant sequence by only one or two single nucleotide polymorphisms (SNPs). Thus, the pre-bottleneck mitochondrial diversity appears only slightly higher than at present and much lower than that observed in older samples. The 14 kyr-old Kesslerloch specimen reveals the first occurrence of the mitogenome lineage of the extant *B. bonasus* population. Thus, the modern population corresponds to a minor fraction of the diversity that was present in Europe during the Late Pleistocene when *B. bonasus* replaced *B. priscus.*

A major reduction in the intrapopulation diversity is apparent from the Late Pleistocene to the early 20^th^ century (Table 1). The Pleistocene populations of Siberia, the Caucasus and Europe are characterized by a high diversity at both the haplotype (H=1.00) and nucleotide levels (as estimated with Pi and various Theta estimators), the nucleotide diversity of the European *B. priscus* being lower than that of its Siberian population. The Western European Holocene population of *B. bonasus* experienced a major reduction of its diversity between the Middle Ages and the beginning of the 20^th^ century.

**Table 1.**

| Population | Nind | Nhap | Haplotype diversity, *H* | Nucleotide diversity, *Pi* | Theta(k) | Theta(S) | Theta(Pi) | Theta (per site) |
| --- | --- | --- | --- | --- | --- | --- | --- | --- |
| Upper Pleistocene Siberia (Bp) | 4 | 4 | 1.00 ± 0.18 | 0.046 ± 0.031 | * | 15.82± 8.85 | 16.83 ± 9.55 | 0.044 |
| Upper Pleistocene Europe (Bp) | 4 | 4 | 1.00 ± 0.18 | 0.014 ± 0.010 | * | 4.91 ± 2.97 | 5.17 ± 3.77 | 0.014 |
| Upper Pleistocene Caucasus (Bb1-Bb2) | 7 | 7 | 1.00 ± 0.08 | 0.078 ± 0.045 | * | 23.67 ± 10.92 | 28.95 ± 16.57 | 0,068 |
| Holocene Europe (ancient, Bb2) | 5 | 3 | 0.80 ± 0.16 | 0.010 ± 0.007 | 2.22 | 4.32 ± 2.48 | 3.80 ± 2.68 | 0,012 |
| Modern Europe (Pre-WWI, Bb2) | 19 | 3 | 0.51 ± 0.12 | 0.001 ± 0.001 | 0.73 | 0.57 ± 0.43 | 0.56 ± 0.53 | 0,0016 |
| Modern Caucasus (Pre-WWI, Bb2) | 4 | 2 | 0.67 ± 0.20 | 0.002 ± 0.002 | 0.88 | 0.54 ± 0.54 | 0.67 ± 0.75 | 0,0015 |

Pairwise Nucleotide divergence with Jukes and Cantor calculated with DNAsp
|  | Upper Pleistocene Siberia | Upper Pleistocene Europe | Upper Pleistocene Caucasus | Holocene Europe (ancient) | Modern Europe | Modern Caucasus |
| --- | --- | --- | --- | --- | --- | --- |
| Upper Pleistocene Siberia | * |  |  |  |  |  |
| Upper Pleistocene Europe | 0.03712 | * |  |  |  |  |
| Upper Pleistocene Caucasus | 0.21127 | 0.19816 | * |  |  |  |
| Holocene Europe (ancient) | 0.19524 | 0.18988 | 0.07614 | * |  |  |
| Modern Europe | 0.18548 | 0.17621 | 0.07498 | 0.01797 | * |  |
| Modern Caucasus | 0.18306 | 0.17443 | 0.07506 | 0.02128 | 0.00367 | * |

Calculations relied only on the samples that yielded the four PCR fragments

We performed a phylogenetic analysis of the HVR under the Bayesian framework estimating mutation and population history parameters from temporally spaced sequence data using all dated ancient and modern bison HVR sequences present in GenBank in 2015 in combination with the data from the current study (Fig. 2). The resulting tree shows a bifurcation between the *Bison bonasus* and *Bison priscus/bison* mitogenome lineages about 1.0 Mya [95% highest posterior density interval (HPD): 1. 5−0.7] (Table 2 and Fig 2). The most recent common ancestor (MRCA) of the Eurasiatic and the American *B. priscus* clades is estimated here at 151 kya [193–119], which is in agreement with a previous estimate of 136 ky [164-111] that was also based on the HVR (7). *B. priscus* from America and Siberia group in distinct clades, and the modern American bison descended from a small subgroup of a once diverse American population, as previously proposed (7). The Late Pleistocene European *B. priscus* corresponds to a subset of the Siberian population.

**Fig. 2.**
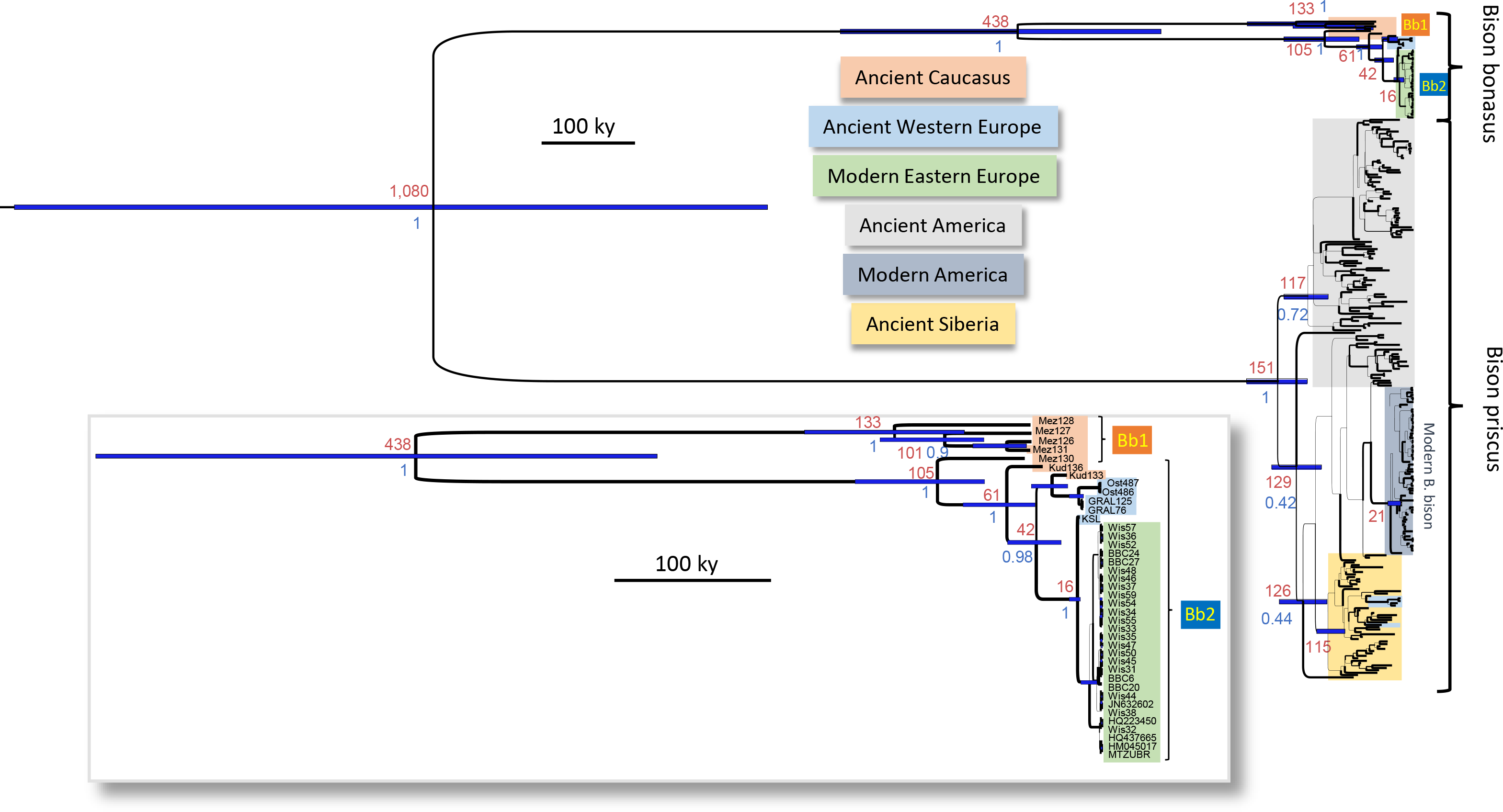
Bayesian phylogeny of Bison hypervariable region. All dated and complete ancient Bison sequences produced here and in a previous *B. priscus* analysis (7) were aligned and reduced to the 367 bp sequence targeted herein alongside modern *B. bonasus* and *B. bison* sequences. A Bayesian phylogenetic analysis was performed using Beast to estimate the age of the nodes from temporally spaced sequence data. The age of the nodes (in kya) is indicated in red, whereas the blue bars represent the 95% HPD of these ages. The color code representing the origin of the various samples is as indicated. The inset on the lower part of the figure represents a magnified view of the *B. bonasus* branches of the tree. The posterior probability of the nodes is indicated in blue and the thickness of the branches is proportional to this posterior probability.

**Table 2.**

| Age estimates of the nodes (kya) |  |  |
| --- | --- | --- |
| Node | Age from HVR [95% HPD] | Age from mitogenome [95% HPD] |
| Root Bovina | 1080 [1550-700] | 944 [1090-810] |
| Bos Taurus/B. bonasus | - | 786 [908-668] |
| B. indicus/B. taurus | - | 161 [189-133] |
| B. grunniens/B. priscus | - | 328 [378-272] |
| B. priscus | 151 [193-119] | 115 [135-95] |
| B. bison | 21 [27-16] | 12 [14-7] |
| All B. bonasus | 438 [643-284] | 250 [287-211] |
| B. bonasus Bb2 | 105 [158-75] | 87 [97-75] |

| Clock rate estimates (per site * year) |  |  |
| --- | --- | --- |
| Partition | Rate from HVR [95% HPD] | Rate from mitogenome [95% HPD] |
| HVR | 5.4 [3.9-6.9] E-7 | 4.6 [3.7-5.6] E-7 |
| RNA | - | 2.5 [2.1-2.9] E-8 |
| Coding (1st-2d positions) | - | 2.6 [2.2-3.0] E-8 |
| Coding (3d position) | - | 9.1 [7.8-10.5] E-8 |

The phylogenetic analyses of the *B. bonasus* haplotypes reveal that the ancient Caucasian population had a deep root and was highly diverse. Based on the HVR, the age of the MRCA of the *Bb1* and *Bb2* clades is estimated at 438 [643–284] kya, which is significantly older than the MRCA of *B. priscus* (Fig. 2 and Table 2). This observation indicates that the Upper Pleistocene Caucasian population had a deeper root than the contemporaneous and more northern *B. priscus* population, even though the latter one may have occupied in the past a much larger territory from Western Europe to the American continent. The *Bb2* clade encompasses several branching events, the separation of a branch represented by a ca. 50 kyr-old northern Caucasus specimen from Mezmaiskaya occurring first, i. e., 105 [158–75] kya, followed, 61 [89–43] kya, by the separation of a branch represented by a ca. 37.8 kyr-old specimen from the Kudaro cave in the southern Caucasus. Then, 42 [60–26] kya, the subgroup comprising an 22.2 ky-old specimen from the Kudaro cave as well as all French specimens between 12.4 to ca. 1.2 kya separated from the haplogroup encompassing the modern *B. bonasus* sequences. Finally, in this later haplogroup, the 14 ky-old Kesslerloch specimen and the early 20^th^ century Central European and Caucasus specimens have a MRCA estimated at 16 [33-14] kya. This sequential order of the radiation events and the previously mentioned reduction of the population diversity indicates that at the transition between the Pleistocene and the Holocene, Western and Central Europe were populated by a subset of a wisent population established in an area of southern Asia including the Caucasus, and that only a part of this population survived into the 20^th^ century.

### The evolution of the Bison clades based on entire mitogenomes

To render this phylogeny and dating scenario more robust, we used biotinylated RNA probes synthesized from the complete mitogenome of *B. p. taurus*, and optimized a sequence capture approach to recover complete mitogenome sequences from seven well-preserved 50 to 12 kyr-old specimens representative of the various clades and radiation events defined above. We performed a Bayesian phylogenetic analysis of these mitogenomes and of two recently published *B. priscus* mitogenomes (14, 23) together with those of modern *B. bison* and *B. bonasus, B. p. taurus*, and *P. grunniens* complete mitogenomes present in GenBank in 2015 (Fig. 3). As previously observed with modern sequences (9, 21, 22), the *B. bonasus* mitogenome lineage is more closely related to the *Bos p. taurus* lineage than to the *B. priscus-B. bison* lineages. The Bayesian analysis reveals, however, that there is a significant overlap (35 to 40%) of the 95% HPD intervals of the dates estimated for the node separating the *Bos p. taurus-B. bonasus* and the *B. priscus-B. bison* lineages, estimated here at 944 [1090-810] kya, and the node separating the *B. p. taurus* and *B. bonasus* lineages, estimated here at 786 [908-668] kya. Such an overlap suggests that the two bifurcation events may have occurred within a relatively short evolutionary period, thus increasing the likelihood that these two events preceded the major separation of the *B. p. taurus* and *Bison* species. This peculiar affiliation pattern of mitogenomes renders the incomplete lineage sorting hypothesis a parsimonious interpretation (see SI Fig. 2). The *Bison priscus/bison* and yak (*P. grunniens*) mitogenome lineages separated from a common ancestor dated at ca. 328 kya [378-272]. For the *Bison* mitogenomes, the dates of the nodes estimated with complete mitogenomes are often younger than those estimated using only the HVR (Table 2). For instance, the common ancestor of the *Bb1* and *Bb2* clades is estimated at 250 kya [287-211] when comparing full mitochondrial genomes, rather than 438 kya [643-284] calculated from only HVR sequences. Similarly, the common ancestor of the Eurasiatic *B. priscus* and the modern American bison is estimated at 115 [135-95] kya rather than 151 [193-119] kya. In contrast, the various branching events within the *Bb2* clade show non-significant differences (given the overlap of the HPD) between the two series of date estimations: 87 [97-75] kya instead of 105 [158-75] kya for the ancestor of the North and South Caucasian specimens, 63 [70-52] kya instead of 61 [89-43] kya for the ancestor of the South Caucasus and Holocene European specimens, and 37 [42-29] kya instead of 42 [60-26] kya for the ancestor of the Holocene European specimens. The discrepancies between the date estimates for several nodes depending on the genetic region used may be partly due to the differences in the number of individual sequences studied in the two series. The major source of discrepancy, however, appears to be due to an irregular rate of evolution of the HVR. Indeed, even though the clock rates estimated for the HVR are similar when either the HVR alone or as a partition of the complete mitogenome is used (respectively 5.4 [3.9−6.9] and 4.6 [3.7−5.6] E-7 per site × year), the individual HVRs show distinct evolutionary rates. When the distribution throughout the mitogenome of the SNPs distinguishing individual bison mitogenomes from the *B. p. taurus* mitogenome as an outgroup are compared, the number of SNPs accumulated in the HVR can vary up to two-fold between individual sequences whereas the rest of the mitogenome is equally distant to the outgroup (SI Fig. 3). For example, there are twice as many mutations accumulated in the HVR subregion (15900-16100) of the Mez 128, Gral 232 and Yaku 118 mitogenomes than in the ones of the Mez130 and Gral 125 specimens. Whatever the underlying mechanism responsible for these differences of the evolutionary rate of this particular region of the mitogenome, this phenomenon limits the reliability of the dating estimations based solely on the HVR.

**Fig. 3.**
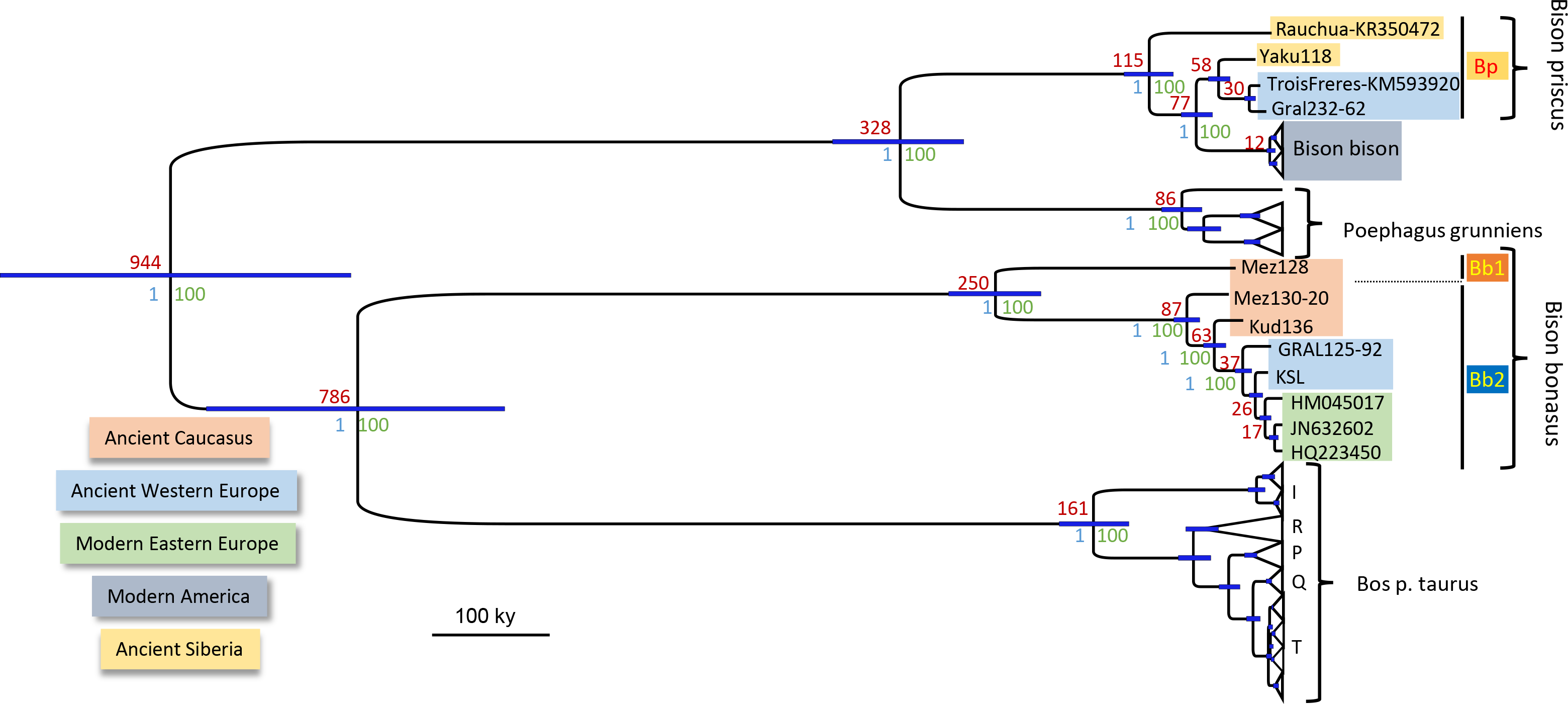
Bayesian phylogeny of complete mitogenomes of *Bos* and *Bison*. We used the complete mitogenomes of ancient *Bison* obtained herein as well as the two published *B. priscus* mitogenomes, and all modern *Bison bison, Bison bonasus, Poephagus grunniens*, and *Bos primigenius taurus* mitogenomes available in Genbank in 2015 totaling 439 sequences. The *B. bison*, *P. grunniens* and *B. p. taurus* sequences have been collapsed to preserve only their subclade structure. The estimate of the age of the nodes, in kya, are indicated in red, with the 95% HPD indicated by blue bars. The statistical supports of the nodes are indicated in blue (Bayesian posterior probability) and in green (bootstrap values of a ML phylogeny performed using RaXML).

Complete mitogenomes confirm the observation made with the HVR sequences that the European *B. priscus* specimens from the Late Pleistocene were more homogeneous genetically than the contemporaneous Siberian population. Indeed, the two French mitogenomes from the Gral (Gral232) and Trois-Frères cave (23) are very similar, with only 19 SNPs distinguishing them, whereas the two Siberian mitogenomes (Yaku118 and Rauchua (14)) are much more divergent with eight-fold more SNPs distinguishing them (153 SNPs). Strikingly, the age of the MRCA of the 26.7 kyr-old Yakutian sample and the 20 to 15 kyr-old French *B. priscus* samples is estimated at 58 [66-47] kya. Since the MRCA of these sequences is also the MRCA of the group of HVR sequences comprising two additional Yakutian and all other French *B. priscus* samples (see Fig. 1), this indicates that the steppe bison inhabiting France between 39 and 15 kya originated from a migration from North-East Eurasia that occurred not earlier than 58 [66-47] kya. Presumably, the low genetic diversity of the Western European *B. priscus* population is due to a founder effect during this migration.

The phylogenetic relationships between the *B. bonasus* mitogenomes indicate that modern wisent in central Europe are more closely related to the Late Pleistocene/Holocene Western European population than to the Late Pleistocene Caucasian population. Indeed, the 14 kyr-old specimen from the site of Kesslerloch at the Swiss-German border is closely related to modern wisent, with only 15 SNPs distinguishing the ancient and modern mitogenome sequences. This demonstrates that the range of the ancestral population of the modern wisent encompassed Middle Europe 14 kya. The Late Pleistocene Caucasian population was highly diverse with a deep root separating most of the specimens from the Mezmaiskaya cave in the northern Caucasus, belonging to the *Bb1* clade, from the specimens from the Kudaro cave in the southern Caucasus, belonging to the *Bb2* clade, and including a specimen from the northern Caucasus: 342 SNPs distinguished the Mez128 (Bb1) and the Mez130 (Bb2) mitogenomes and 105 SNPs distinguished the northern (Mez 130) and southern Caucasian (Kud136) mitogenomes of the *Bb2* clade. In contrast, the members of the *Bb2* clade in Western Europe were more similar, with only 39 SNPs distinguishing the French (Gral125) and Swiss (KSL) specimens, in agreement with the reduction of diversity that we observed when comparing the HVR of the Western European and Caucasian *B. bonasus* populations. Thus, complete mitogenomes confirm the observations from the HVR analyses of a larger sample size while providing a more accurate phylogenetic analysis, in particular with respect to the dating of the MRCA of the various mitogenomes.

## Discussion

### European bison populations turnover during the late Pleistocene and the Holocene

Our results enable us to propose a scenario for the evolutionary history of the bison in Europe that is related to the climatic fluctuations and the resulting environmental changes of the Late Pleistocene and the Pleistocene/Holocene transition as summarized in Figure 4. We observe striking regional and temporal differences in the major clades and distinguish three periods, particularly in France. The first period, from at least 47 kya to about 34 kya, was characterized by the dominance of a divergent *B. bonasus* lineage belonging to the *Bb1* clade in both southern France (Arquet and Plumettes, 7 *Bb1* out of 7 samples) and the northern Caucasus (Mezmaiskaya, 5 *Bb1* and 1 *Bb2* out of 6 samples). This lineage was absent from the samples from later periods indicating that the corresponding population was the first to disappear. For the same time period, the steppe bison mitotype *Bp* was the sole mitotype found in Siberia and northern Eurasia (27/27 samples dated from 66 to 34 kya (7)). This period, encompassing most of MIS3, is characterized by oscillating shorter glacial and longer interstadial periods, the latter lasting more than one thousand years (glacial interstadial GI15 to GI8 (12, 25) and Fig. 4). During the warmer periods, southern France and the northern Mediterranean coast were covered by deciduous coniferous non-continuous forests, and central France and central Europe by coniferous open woodland (26). The steppe bison *B. priscus* was dominant in France during the following period that lasted until the end of the Last Glacial Maximum (LGM) at 14.7 kya (7 out of 7 samples, plus 1 out of 1 in (23)). These bison were closely related to the Siberian sample from Yakutia from the same period, the whole mitogenome of which we analyzed (99.4% identity between the Yaku128 and Gral232 mitogenomes). HVR comparisons between the French and Siberian and North American samples (7) reveal that the French *B. priscus* mitotypes are closely related to those of the Siberian samples (Fig. 2), indicating that the Western European territory was colonized by a subpopulation of the steppe bison from the northern Eurasian continent. The age estimated from full mitogenomes for the MRCA of the French and Siberian samples is 56 [64-46] kya, indicating that this colonization involved a population that separated from the Siberian population more recently than ca. 56 kya and suggesting a lower limit for the arrival in France of these steppe bison. In contrast, specimens older than 60 kya assigned to *B. priscus* in the fossil record of Western Europe presumably must have belonged to a distinct population that was not the direct ancestor of this *B. priscus* population that occupied Western Europe during the cold spells of MIS2. At the end of MIS3, around 32 kya, the climate became colder on average and the warmer interstadials were shorter, lasting only a few hundred years. Then, between 27 and 14.7 kya, a second, long glacial period followed that comprised two phases. In the first phase, the tree cover was patchy and incomplete, with a high proportion of steppe vegetation, whereas the second, a full glacial phase, was characterized by sparse grassland and open steppe tundra in southern and northern Europe, respectively (27). While during this period the steppe bison *B. priscus* occupied the territory previously occupied by *B. bonasus* in Western Europe, *B. bonasus* remained nevertheless present in the southern Caucasus, even during the LGM. Indeed, in our samples, two out of two specimens from the southern Caucasus belonged to the *Bb2* clade, which we found to be present at low frequency at an earlier period in the northern Caucasus. Finally, during the third period, starting at the end of the MIS2 and lasting up to the present, *B. bonasus* of the *Bb2* clade expanded again into Western Europe, as we detected it in the 14 kyr-old specimen from Switzerland at the beginning of the Bølling-Allerød interstadial period (14.7 to 12.7 kya). The more recent specimens from France belonged without exception to the *Bb2* clade (7 out of 7, dated between 12 kya to the Middle Ages). Strikingly, the sedimentary sequence of the French site Gral recorded a population replacement: all specimens older than ca. 15 kya, before the B0lling-Aller0d interstadial, belong to the *B. priscus Bp* mitotype (5 out of 5), whereas all more recent ones, coinciding with the onset of a more temperate climate at the end of the last glacial event (end of MIS2), belong to the *B. bonasus Bb2* mitotype (2 out of 2, Pval<0.05 with Fisher’s test). The ancient French *Bb2* population is significantly different from the one comprising the modern wisent and the ancient Kesslerloch specimen (Fig. 1-3). Moreover, this subclade was not detected later than the Middle Ages, suggesting that it went extinct in Western Europe with the disappearing local wisent. In contrast, the distinct *Bb2* mitotype of the 14 kyr-old sample from Kesslerloch continued to exist up to present time. It is the only remaining mitotype detected in both present-day wisent as well as in the specimens from Poland and the Caucasus from the beginning of the 20^th^ century prior to the last major bottleneck of WWI.

The marked climatic variations that occurred during the Pleistocene implied drastic environmental changes that triggered the profound reorganization of floral and faunal biomes at different geographic and temporal scales. Response of biota to climatic stimuli was regionally/continentally distinct, sometimes occurring synchronously but most often diachronously (e.g., (2, 28)). The repeated cyclical climate changes progressively promoted landscape renewal, locally modifying, obliterating and creating specific and varied ecological niches (28). The various expansions and contractions of bison populations in response to these climatic fluctuations led to apparent alternating occupations of Western Europe by either *B. priscus* or *B. bonasus* during the last 50 kyrs, each presumably originating respectively from either a northeastern territory including Siberia, or a southern territory including the Caucasus. We hypothesize that these alternating occurrences result from climate-driven changes of two different habitats to which *B. priscus*, a grazer, and *B. bonasus*, a mixed feeder, were adapted, i.e., the open tundra-steppe for the first and open woodlands for the second. Indeed, the diet of *B. priscus* during the LGM included typical steppe and grassland (C_3_) vegetation and lichens (29), whereas the wisent’s diet in the Holocene was more flexible and included a higher content of shrubs (18). Similarly, on the American continent, *B. priscus* adapted to the climatic and environmental changes of the Holocene and evolved in two recently divergent forms, the plain bison thriving on the grasslands of the Great Plains and the wood bison inhabiting the boreal forest in North America. This suggests that on the Eurasian continent the competition with *B. bonasus*, a species seemingly better adapter to a more temperate environment, may have prevented a similar adaptation of *B. priscus* to habitat changes during the warmer periods. Finally, in Western Europe, local variations in ecological and physical barriers could have affected the speed of bison population turnover as a reaction to climatic shifts. In the future, a higher resolution genetic study involving a much higher sample number with denser time and space sampling may provide a more accurate and nuanced view of these population turnovers.

**Fig. 4.**
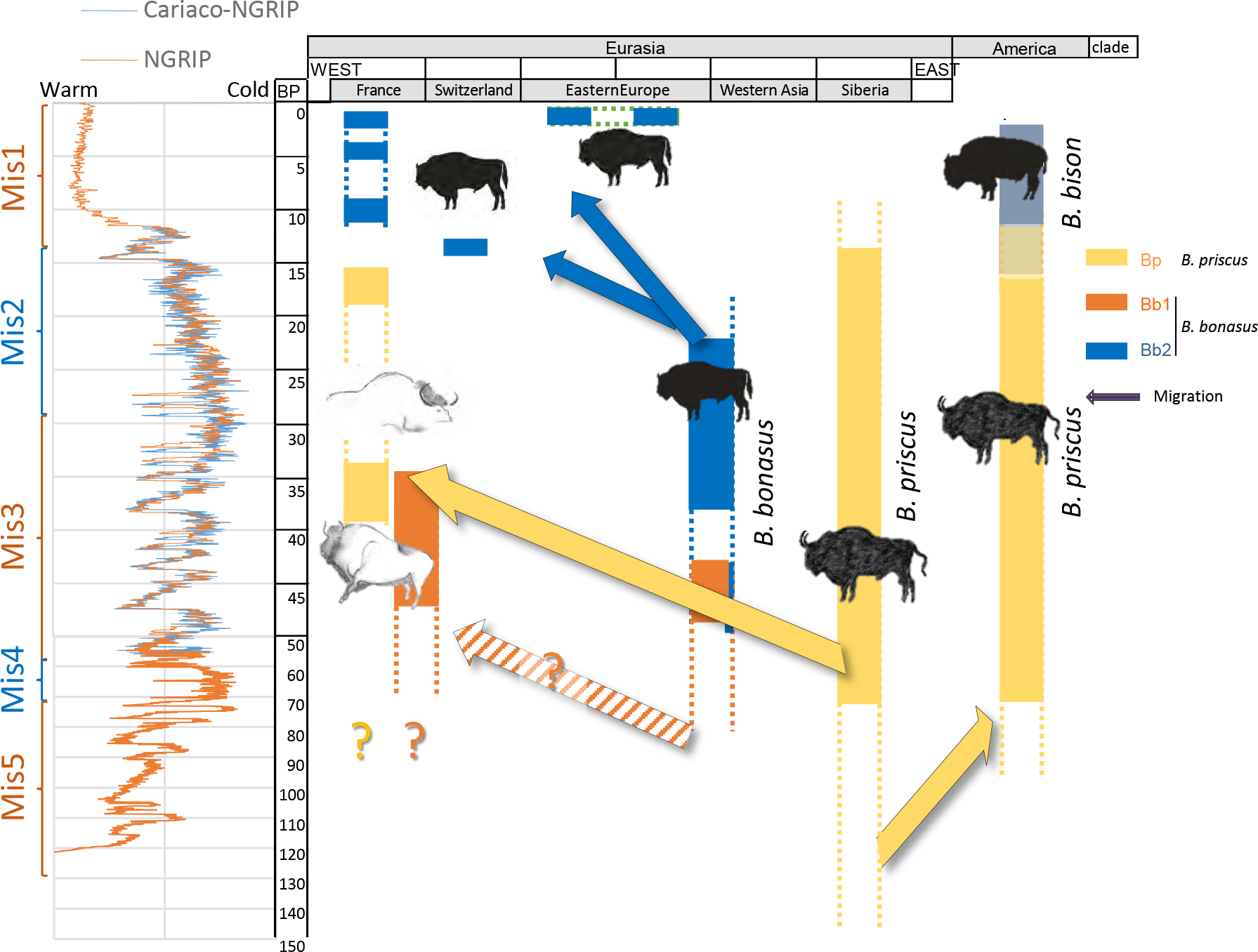
Schematic representation of the distribution through time and space of the various mitogenome clades. The geographic regions are represented on the abscissa and the time scale on the ordinate. The ascertained presence of the various mitochondrial haplogroups are represented by solid boxes, whereas the dotted lines indicate possible temporal extension of the presence of these clades. The left side shows the climatic fluctuations as inferred from the North Greenland Ice Core Project (NGRIP) (25) and the combined Carabean Cariaco basin and NGRIP data as shown in (12), as well as the Marine Isotope Stage (MIS) as defined by Lisiecki and Raymo ^24^ (http://www.lorraine-lisiecki.com/LR04_MISboundaries.txt). The proposed migrations are indicated by solid arrows. The hatched arrow indicates a possible migration of the *Bb1* clade that populated Western Europe from a southern refugee before the time period analyzed herein. The genetic identity of the bison that, according to the fossil record, populated Western Europe before 60 kya is not known, but climatic fluctuations may have triggered additional expansions and contractions of different populations of *B. priscus* and *B. bonasus.* (Drawings: E.-M. Geigl)

The mitochondrial lineages of *B. bonasus* that were present 40 kya have an older root than those of *B. priscus* (250 [287-211] vs 115 [135-95] kya). This indicates that the *B. bonasus* did not experience as severe a bottleneck prior to 100 kya as the population reduction the steppe Bison experienced between 150 and 100 kya, before later thriving in northern Eurasia, Beringia, and in North America between 80 and 20 kya. All the various population expansions and contractions that occurred in response to the climatic and environmental changes characterizing the late Pleistocene and the transition to the Holocene gave rise to a reduction of the population diversity and to the extinction of lineages, like the *Bb1* lineages, which apparently was the first to disappeared, and the *Bp* lineage, which disappeared at the beginning of the Holocene on the Eurasian continent. Finally, the reduction of the diversity of the *Bb2* lineage seems to involve both migrations and local extinctions with the major and most severe reductions occurring between the Early Holocene and historic times, most likely owing to human impact through hunting pressure and habitat fragmentation.

Numerous cave art representation in southern France and northern Spain, and also in the Caucasus suggest realistic depictions of both the steppe bison and the wisent (19). We consider likely that the paintings of bison, the so-called “bison of the pillar”, in the cave Chauvet-Pont d’Arc in France depict the two types of bison distinguished by the shape of horns and back lines (Fig. 5). The upper image on the pillar, dated to 38.5-34.1 kya (30), could represent a wisent, and the lower image, dated to 36.2 to 34.6 kya (30), a steppe bison. These dates coincide with the period (39 to 34 kya) in which the two bison forms overlap in our dataset from southern France, in the vicinity of these paintings. Within this time frame, wisent population would have been in decline and steppe bison population would have been expanding.

**Fig. 5.**
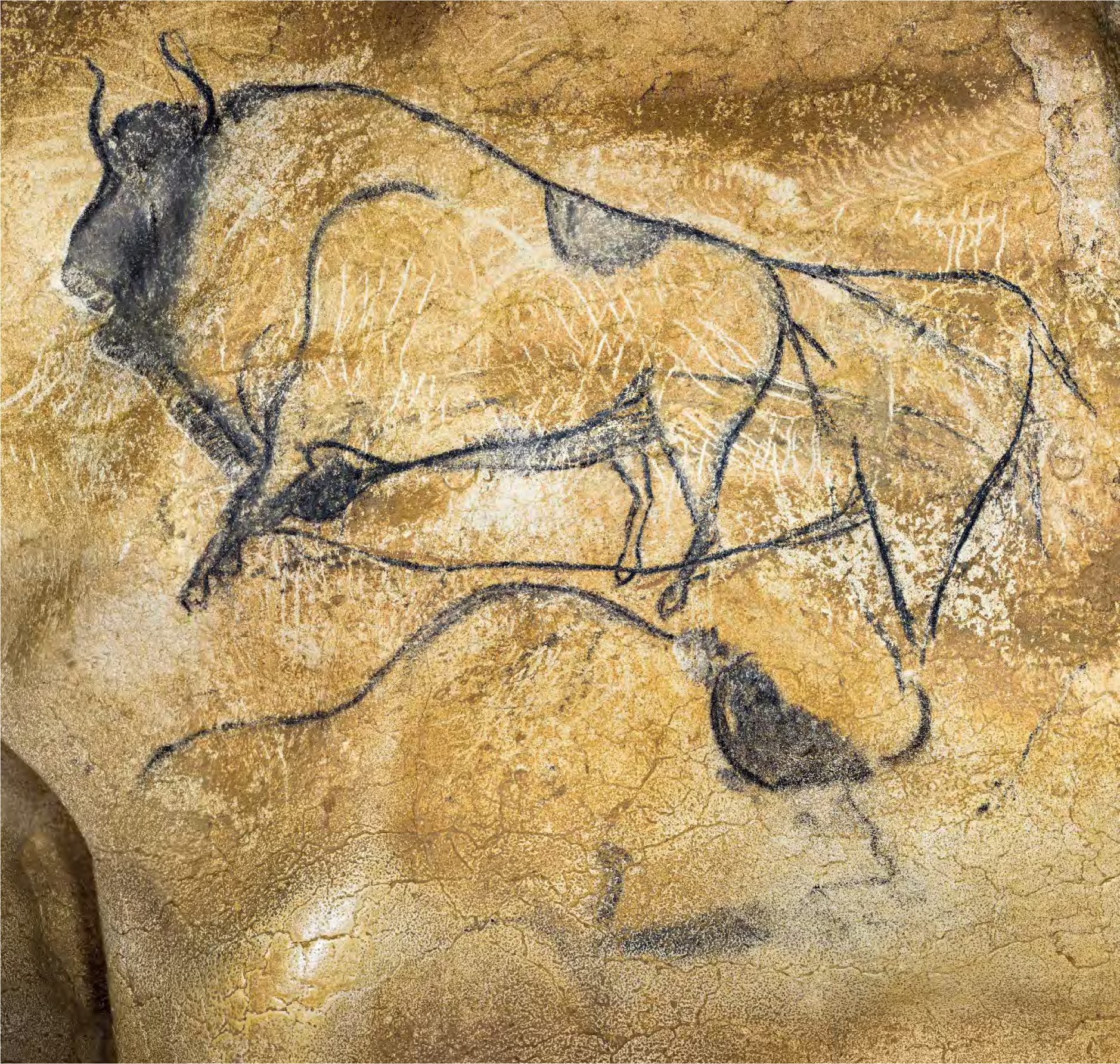
Prehistoric painting of bison in the cave of Chauvet-Pont d’Arc, Ardeche, France. The paintings are the so-called “Bison of the pillar” in the “End Chamber” of the Chauvet cave. The charcoal of both drawings have been radiocarbon dated at 38.5-34.1 kya for the upper bison, and at 36.3-34.6 kya for the lower bison (30). We consider, based on criteria stated by Spassov (19) that the “great bison” in the upper part represents *B. bonasus* with a highly positioned head, curved horns, a moderately large hump and a weak mane and rather equilibrated body proportions between the front and the rear. The lower part would represent *B. priscus* with its large hump, its low head position, its abundant mane, crescent-shaped horns, and, although somewhat faded in the image, the steep incline of the back-line and stronger hindquarters can be made out. (Copyright: French Ministry of culture, archeologie.culture.fr/chauvet. Cl. Arnaud Frich. CNP/MCC)

### Radiation of the *Bos* and *Bison* lineages

Since the initial classification of Linnaeus in 1758 of *Bos* and *Bison* within a single genus, there has been debate about whether they should be placed in separate genera (31). Indeed, the members of these genera can still be crossed, despite reduced fertility in some combinations of crosses, and *B. p.taurus* genetic material can be found in a number of present-day American bison and wisent (32, 33). Thus, gene flow could have occurred between the various lineages, in particular in the early phases of their differentiation. Such gene flow between separated populations has already been considered as an alternative possibility to incomplete lineage sorting to explain that the mitogenome of *B p. taurus* was closer to that of *B. bonasus* than to that of *B. bison* (21, 22). The phylogenetic estimates of the age of the various common ancestors of these two lineages indicate, however, that the most parsimonious interpretation of incomplete lineage sorting suffices to account for the relative affiliations of these mitogenomes (see SI Fig. 2). Indeed, as mentioned above, the age estimates indicate that the MRCA of the *Bos* and *Bison* mitogenome lineages is not much older than that of the *B. p. taurus* and *B. bonasus* lineages, respectively 946 [1090-810] kya, and 787 [908-670] kya, with a 35 to 40% overlap of the confidence interval (95% HPD). This indicates that the two bifurcation events have occurred within a relatively short evolutionary period. Rapid speciation during this short evolutionary period appears unlikely since these species have still not yet totally lost interfertility almost a million years later. Our radiation date estimates are within the same range of the earliest fossils that were clearly attributed to the *Bos* genus and that are dated at 1 Mya (34). They are also consistent with the analysis of the complete genome of a modern wisent that estimated that the wisent and bovine species diverged between 1.7 to 0.85 Mya through a speciation process involving an extended period of limited gene flow with some more recent secondary contacts posterior to 150 kya (35). Thus, incomplete lineage sorting of mitogenomes in a metapopulation of the *Bos* and *Bison* ancestors during the period of divergence of these species could account for the affiliation patterns of these mitogenomes without the need to postulate a more recent post-speciation gene flow. It is interesting to note that this radiation event could be coincident with the onset and intensification of high-latitude glacial cycles (100 kyr-periodicity) around 1.2-0.8 Mya. Incomplete lineage sorting, however, does not preclude later sporadic introgression of nuclear DNA at any point up to the present owing to the persistent interfertility between these species, as evidenced by the detection in the wisent genome of ancient gene flow from *B. p. taurus* (35, 36).

## Conclusion

The analysis of DNA preserved in ancient bison remains from Eurasia covering the last 50,000 years allowed us to retrace some of the population dynamics that took place during the Late Pleistocene and the Holocene, including population migrations, extinctions and replacements. We could trace back the origin of the wisent to Eastern Europe including the Caucasus. We observed several alternating expansion waves of wisent and steppe bison in Western Europe that were related with climatic fluctuations, the wisent being prevalent when the climate was more temperate leading to a more forested vegetation whereas the steppe bison population originating from northern Eurasia was predominant in Western Europe during the colder periods of the Late Pleistocene with their open environments. These fluctuations may have been recorded in Paleolithic cave paintings, in particular in the cave of Chauvet that had been occupied by humans over a long period and where two distinct bison types are depicted.

## Materials & Methods

Details of samples are described in the Supplementary Information (SI Table 1 and SI Fig. 4). All pre-PCR procedures were carried out in the high containment facility of the Jacques Monod Institute physically separated from areas where modern samples are analyzed and dedicated exclusively to ancient DNA analysis using the strict procedures for contamination prevention previously described (37).

### DNA Extraction

The external surface of the specimens was removed with a sterile blade to minimize the environmental contamination. For each bone sample, roughly 0.2 g was ground to a fine powder in a freezer mill (Freezer Mill 6750, Spex Certiprep, Metuchen, NJ), which was then suspended in 2mL of extraction buffer containing 0.5M EDTA, 0.25M Di-sodium Hydrogen Phosphate (Na_2_HPO_4_), pH 8.0, 0.14M 2-mercaptoethanol and 0.25 mg/mL of proteinase K and incubated under agitation at 37°C for 48 hours. Blank extractions were carried out for each extraction series, eight in total.

Samples were then centrifuged and the supernatant was purified with the Qiaquick PCR Purification Kit (Qiagen, Hilden, Germany), as described (37).

### PCR Amplification of the HVR

Primers were designed to target conserved regions with minimal primer dimer propensity against a multiple alignment of all the Bison mitochondrial sequences present in Genbank in 2012 using the software Oligo 7 as described (38, 39), and were then tested for efficiency and dimer formation using quantitative real-time PCR (qPCR) (38, 39). Four primer pairs amplifying short (118 to 152 bp) subregions of the mitochondrial Hyper Variable Region (HVR) were selected (SI Table 2 and SI Fig. 1). To protect against cross-contamination between samples, we used the UNG-coupled quantitative realtime PCR system (40) and, to avoid the production of erroneous data due to the presence of bovine DNA in reagents, we decontaminated reagents as previously described (41). Amplifications were performed in a final volume of 10 μL containing 2mM MgCl_2_, 1μM primers, 0.04mM of dA/G/CTPs and 0. 08mM of dUTP, 0,01U/μL of UNG, 1U/μL of FastStart Taq (Roche Applied Science) and 1 × qPCR home-made reaction buffer (41). Blank amplification controls were included for each amplification. In total, 415 amplification blanks were carried out during the various amplifications analyses of 85 samples. No products were observed in any of the amplification blanks. Amplifications were performed using a LightCycler 1.5 (Roche Applied Science) using the following cycling program:15 minutes at 37°C (carry-over contamination prevention through digestion by UNG of dUTP-containing amplicons), 10 minutes at 95°C (inactivation of UNG and activation of the Fast Start DNA polymerase) followed by 60 cycles at 95°C for 15 seconds and 60°C for 40 seconds (for primer pairs Bon1, 2 and 3) and 95°C for 15 seconds, or 56°C for 15 seconds and 67°C for 20 seconds (for primer pair BB3r4m), followed by a final melting curve analysis step. All extracts were tested for inhibition as previously described (42). Each sequence was determined on both strands from at least two independent amplifications using capillary electrophoresis sequencing. Sequencing data analyses were performed using the software Geneious 6.1.8 (43).

### Capture and sequencing of whole mitogenomes

Dual barcoded libraries for Illumina sequencing were constructed using a double-stranded procedure previously described ((37, 44), see also Supplementary Methods). The End-repair and adapter ligation mixtures were decontaminated with Ethidium monoazide (45) to inactivate bovine DNA associated with BSA present in these reagents that could interfere with the final results. For the production of complete mitogenome sequences, we developed a sequence capture approach using biotinylated RNA baits (46), further detailed in the supplementary information. Briefly, we amplified the *Bos taurus* mitogenome by PCR targeting eleven 1.5 kb-long fragments using primer pairs, one of which contained the T3 promoter as a 5’extension. Strand-specific biotinylated RNA probes were synthesized, pooled and submitted to mild, controlled heat-induced hydrolysis to generate RNA fragments ranging from 100 to 600 nucleotides with an average size of 300. Each library was individually hybridized to the biotinylated probes in the presence of RNA competitors complementary to the Illumina adapters for 48 hours at 62°C. After three 10-min 0.1X SSC washes at 62°C, captured DNA was eluted by alkaline hydrolysis of the RNA probes, amplified an optimal number of PCR cycles as predetermined by qPCR, and subjected to a second round of sequence capture in identical conditions. The eluted DNA was then amplified minimally, purified and sequenced on a MiSeq sequencer (Illumina) using paired-end sequencing for 2×75 cycles.

Paired-end reads were merged using leeHom using the—ancientdna parameter (47). Merged reads were then mapped to modern *Bison bison* and *Bison bonasus* mitogenomes using bwa as described in (37). The first 100 nt of each linearized reference mitogenome used for mapping was duplicated at the 3′end to allow mapping of fragments overlapping the junction. Mapped read duplicates were then removed using samtools rmdup as described in (37). The resulting bam files were then imported into Geneious 6.1.8 (43) and remapped onto the appropriate mitogenome sequences without the 100 nt duplication (B. *bison* for *B. priscus* sequences, *B. bonasus* for the Bb1 and Bb2 sequences). Consensus sequences were generated in Geneious and verified by visual inspection of the aligned reads.

### Phylogenetic analyses

Sequence alignments were performed using the Muscle algorithm and were visually inspected and adjusted using Geneious 6.1.8 (43) The maximum likelihood (ML) analyses presented in Fig. 1 were computed using PHYML 3.0, using an HKY substitution model with a gamma-distributed rate of variation among sites (+G) and invariant sites (+I) (48). Robustness of the nodes was estimated using 500 bootstraps. RaXML 8.2.3 was used to generate the ML bootstrap support values for the complete mitogenome alignment shown in Fig. 3 (49).

Phylogenetic analyses conducted under the Bayesian framework were performed using the program BEAST v. 1.8.2, which allows estimation of mutation and population history parameters simultaneously from temporally spaced sequence data (50). Nucleotide substitution models were chosen following comparisons performed with jModelTest 2.1.7 using the Bayesian Information Criteria (51). The HVR analysis presented in Fig. 2 was performed considering a TN93 model for the nucleotide substitution model, a gamma-distributed rate of variation among sites (+G) with four rate categories and invariant sites (i.e., TN93+I+G model). For the complete mitogenome analysis presented in Fig. 3, we used four partitions, the HVR, the 1^st^ and 2^nd^ positions of the codons within the coding region, the 3^rd^ position, and the RNA genes. We considered the HKY+I+G model for the first two partitions, and the TN93+G for the last two. Default priors were used for all parameters of the nucleotide substitution model. For the analysis of figure 2, we used a strict molecular clock with a lognormal prior for the substitution rate (mean=−15.0, stdev=1.4) corresponding to a median of 2E-7 substitutions per site and per year [95%HPD 1.3E-8, 3.2E-6] based on the estimation for Bison HVR substitution rate (7). For the various partitions of the mitogenome of Fig. 3, we used estimates for the human mitogenome substitution rate to set the priors (52): HVR, mean=−16.1 stdev=2.0, corresponding to a median of 1.0E-7 [2E-9-5E-6]; RNA, mean=−18.65 stdev=2.0, corresponding to a median of 8.0E-9 [1.6E-10-4E-7]; 1^st^ & 2^nd^ positions, mean=−18.5 stdev=2.0, corresponding to a median of 9.0E-9 [1.8E-10-4.7E-7]; 3^rd^ position, mean=−17.7 stdev=2.0 corresponding to a median of 2.0E-8 [4E-10-1E-6]. Finally, a standard coalescent model was considered for the tree prior with a Bayesian skyline plot to model populations (5 and 10 populations with default parameters for the HVR and the mitogenomes respectively). The prior for the tree height followed a log-normal distribution, mean=14.6 stdev=0.6, truncate to 8.0E6 and 1.0E5, corresponding to a median of 2.2E6 and a 95%HPD of [6.3E6-6.7E5], which integrates the various fossil finds assumed to correspond to ancestors of cattle and bison (53, 54).

To estimate the posterior distribution of each parameter of interest, we used the Markov Chain Monte Carlo algorithm implemented in the BEAST software. We ran five independent chains with initial values sampled as described above and an input UPGMA tree constructed using a Juke-Cantor distance matrix. Each of these chains was run for 50,000,000 iterations and for each parameter of interest, 18,0 samples (one every 2,500 generated ones) were drawn after discarding a 10% burn-in period. The BEAST output was analyzed with the software Tracer v. 1.6 http://tree.bio.ed.ac.uk/software/tracer/. Visual inspection of the traces and the estimated posterior distributions suggested that each MCMC had converged on its stationary distribution. Using Logcombiner v. 1.8.2, we further combined all the results from the 5 independent chains. The maximum clade credibility tree with the median height of the nodes was finally calculated using TreeAnnotator v. 1.8.2 and visualized using FigTree v. 1.4.2 http://tree.bio.ed.ac.uk/software/figtree/.

## Acknowledgments

The study was supported by the French national research center CNRS. The paleogenomic facility obtained support from the University Paris Diderot within the program “Actions de recherche structurantes”. The sequencing facility of the Institut Jacques Monod, Paris, is supported by grants from the University Paris Diderot, the Fondation pour la Recherche Médicale (DGE20111123014), and the Région Ile-de-France (11015901). We thank Lou Saïer for the production of HVR data for some samples. We acknowledge the contribution to sampling of Myriam Boudadi-Maligne, Jean-Baptiste Mallye, Pierre Pétrequin, Sylvie Lourdaux, and Rene Remond. We thank G. Baldacci for continuous support. M. T. also acknowledges financial support through the “Biodiversity of East-European and Siberian large mammals on the level of genetic variation of populations (BIOGEAST)” project within the 7th Framework Programme, Marie Curie Actions, that allowed sampling to take place in Russia. We like to thank Marie-Hélène Moncel and Marie-Anne Julien for providing samples that did not yield genetic results, and the French ministry of culture for providing the image of the wall paintings of the Chauvet cave.

## Author Contributions

EMG and TG designed and supervised the overall research with an initial input from MT. DM developed sequence capture and produced the mitogenome data. SG produced the HVR data. TG, EMG, DM and SG analyzed and interpreted the data. TG, EMG and EAB wrote the manuscript with inputs from DM and SG. JPB provided samples, discussed the data and corrected the manuscript. MT, RMA, GBa, GBo, JCC, SM, OP, NS and HPU provided samples and feedbacks on the manuscript.

## Bibliography

1. Hewitt GM (2004) Genetic consequences of climatic oscillations in the Quaternary. Phil. Trans. RL. Soc. Lond. B 359:183–195.

2. Markova AK, et al. (2015) Changes in the Eurasian distribution of the musk ox (Ovibos moschatus) and the extinct bison (Bison priscus) during the last 50 ka BP. Quatern Int 378:99-110.

3. Guthrie RD (1990) Frozen fauna of the Mammoth Steppe. The story of Blue Babe. (The University of Chicago Press, Chicago and London) p 323.

4. Brugal JP (1994-1995) Le Bison (Bovinae, Artiodactyla) du gisement Pleistocene moyen ancien de Durfort (Gard, France). Bull.Mus.nat.Hist.Nat., Paris 4e ser. 16, sect.C, n°2–4:349– 381.

5. Brugal JP, David F, Enloe J, & Jaubert J (1999) Le Bison, gibier et moyen de subsistance des hommes du Paleolithique aux Paleoindiens des grandes plaines (Antibes).

6. Flerow CC (1979) Systematics and Evolution. European Bison-Morphology, systematics, evolution, ecology (in Russian), ed Sokolov VE (Nauka (USSR Ac. of Sc.), Moscou), pp 9–127

7. Shapiro B, et al. (2004) Rise and fall of the Beringian steppe bison. Science 306(5701):1561– 1565.

8. Groves CP (1981) Systematic relationship in the Bovini (Artiodactyla, Bovidae). Z.Zool.Syst.Evolutionforsch. 19(4):264–278.

9. Hassanin A, An J, Ropiquet A, Nguyen TT, & Couloux A (2013) Combining multiple autosomal introns for studying shallow phylogeny and taxonomy of Laurasiatherian mammals: Application to the tribe Bovini (Cetartiodactyla, Bovidae). Molecular phylogenetics and evolution 66(3):766–775.

10. Kowalski K (1967) The evolution and fossil remains of the European Bison. Acta Theriologica 12(21):335–338.

11. Krasinska M & Krasinski ZA (2013) European Bison: The Nature Monograph. 2^nd^ Edition (Springer, Heidelberg).

12. Cooper A, et al. (2015) PALEOECOLOGY. Abrupt warming events drove Late Pleistocene Holarctic megafaunal turnover. Science 349(6248):602–606.

13. Boeskorov GG, et al. (2014) Preliminary analyses of the frozen mummies of mammoth (Mammuthus primigenius), bison (Bison priscus) and horse (Equus sp.) from the Yana-Indigirka Lowland, Yakutia, Russia. Integrative zoology 9(4):471–480.

14. Kirillova IV, et al. (2015) An ancient bison from the mouth of the Rauchua River (Chukotka, Russia). Quatern. Res. 84(2):232–245.

15. Degerbol M & Iversen J (1945) The Bison in Denmark. A zoological and geological investigation of the finds in Danish Pleistocene deposits (Danm.Geol.Unders0gelse, Copenhagen).

16. Geist V (1971) The relation of social evolution and dispersal in ungulates during the Pleistocene, with emphasis on the old world Deer and the genus Bison. Quaternary Research 1(3):283–221.

17. Rautian GS, Kalabushkin BA, & Nemtsev AS (2000) A New Subspecies of the European Bison, Bison bonasus montanus ssp. nov. (Bovidae, Artiodactyla). Doklady Biological Sciences 375 636–640.

18. Bocherens H, Hofman-Kaminska E, Drucker DG, Schmolcke U, & Kowalczyk R (2015) European bison as a refugee species? Evidence from isotopic data on Early Holocene bison and other large herbivores in northern Europe. PLoS One 10(2):e0115090.

19. Spassov N & Stovtechev T (2003) On the origin of the wisent, Bison bonasus (Linnaeus, 1758): presence of the wisent in the upper Palaeolithic art of Eurasia. Advances in Vertebrate Paleontology « Hen to Panta »:125–130.

20. Potter BA, et al. (2010) History of Bison in North America. American Bison: Status survey and conservation guidelines 2010, eds Gates CC, Freese CH, Gogan PJP, & Kotzman M (IUCN, Gland, Switzerland), pp 13–18.

21. Verkaar EL, Nijman IJ, Beeke M, Hanekamp E, & Lenstra JA (2004) Maternal and paternal lineages in cross-breeding bovine species. Has wisent a hybrid origin? Mol Biol Evol 21(7):1165–1170.

22. Zeyland J, et al. (2012) Tracking of wisent-bison-yak mitochondrial evolution. Journal of applied genetics 53(3):317–322.

23. Marsolier-Kergoat MC, et al. (2015) Hunting the Extinct Steppe Bison (Bison priscus) Mitochondrial Genome in the Trois-Freres Paleolithic Painted Cave. PLoS One 10(6):e0128267.

24. Lisiecki LE & Raymo ME (2005) A Pliocene-Pleistocene stack of 57 globally distributed benthic 518O records. Paleoceanography 20(1):1–17.

25. Andersen KK, et al. (2004) High-resolution record of Northern Hemisphere climate extending into the last interglacial period. Nature 431(7005):147–151.

26. Van Andel TH & Tzedakis PC (1996) Palaeolithic landscapes of Europe and environs, 150,000 25,0 years ago: An overview. Quat Sci Rev 15(5–6):481–500.

27. Guiot J, Pons A, de Beaulieu JL, & Reille M (1989) A 140,000 year climatic reconstruction from two European pollen records. Nature 338: 309–313.

28. Palombo MR (2014) Deconstructing mammal dispersals and faunal dynamics in SW Europe during the Quaternary. Quat Sci Rev 96:50–71.

29. Julien M-F, et al. (2012) Were European steppe bison migratory? 18O, 13C and Sr intra-tooth isotopic variations applied to a pallaeoethological reconstruction. Quarternary International 271:106–119.

30. Quiles A, et al. (2016) A high-precision chronological model for the decorated Upper Paleolithic cave of Chauvet-Pont d’Arc, Ardeche, France. Proc Natl Acad Sci U S A 113(17):4670–4675.

31. Boyd DP, Wilson GA, & Gates CC (2010) Taxonomy and nomenclature. American Bison: Status survey and conservation guidelines 2010, eds Gates CC, Freese CH, Gogan PJP, & Kotzman M (IUCN, Gland, Switzerland), pp 13–18.

32. Yudin NS, et al. (2012) Detection of mitochondrial DNA from domestic cattle in European bison (Bison bonasus) from the Altai Republic in Russia. Anim Genet 43(3):362.

33. Halbert ND & Derr JN (2007) A comprehensive evaluation of cattle introgression into US federal bison herds. The Journal of heredity 98(1):1–12.

34. Martfnez-Navarro B, Rook L, Papini M, & Libsekal Y (2010) A new species of bull from the Early Pleistocene paleoanthropological site of Buia (Eritrea): Parallelism on the dispersal of the genus Bos and the Acheulian culture. Quatern Int 212(2):169–175.

35. Gautier M, et al. (2016) Deciphering the wisent demographic and adaptive histories from individual whole-genome sequences. Mol Biol Evol In press, biorxiv http://dx.doi.org/10.1101/058446.

36. Wecek K, et al. (2016) Complex admixture preceded and followed the extinction of wisent in the wild. bioRxiv 059527:doi: http://dx.doi.org/10.1101/059527

37. Bennett EA, et al. (2014) Library construction for ancient genomics: single strand or double strand? BioTechniques 56(6):289–290, 292-286, 298, passim.

38. Cote NM, et al. (2016) A New High-Throughput Approach to Genotype Ancient Human Gastrointestinal Parasites. PLoS One 11(1):e0146230.

39. Guimaraes S, et al. (submitted) A cost-effective high-throughput metabarcoding approach powerful enough to genotype ~44,000 year-old rodent remains from Northern Africa. Molecular Ecology Resources.

40. Pruvost M, Grange T, & Geigl E-M (2005) Minimizing DNA contamination by using UNG-coupled quantitative real-time PCR on degraded DNA samples: application to ancient DNA studies. BioTechniques 38(4):569–575.

41. Champlot S, et al. (2010) An efficient multistrategy DNA decontamination procedure of PCR reagents for hypersensitive PCR applications. PLoS ONE 5(9).

42. Pruvost M, et al. (2007) Freshly excavated fossil bones are best for amplification of ancient DNA. Proc. Natl. Acad. Sci. USA 104(3):739–744.

43. Kearse M, et al. (2012) Geneious Basic: an integrated and extendable desktop software platform for the organization and analysis of sequence data. Bioinformatics 28(12):1647– 1649.

44. Gorge O, et al. (2016) Analysis of Ancient DNA in Microbial Ecology. Methods in molecular biology 1399:289–315.

45. Rueckert A & Morgan HW (2007) Removal of contaminating DNA from polymerase chain reaction using ethidium monoazide. Journal of microbiological methods 68(3):596–600.

46. Gnirke A, et al. (2009) Solution hybrid selection with ultra-long oligonucleotides for massively parallel targeted sequencing. Nature biotechnology 27(2):182–189.

47. Renaud G, Stenzel U, & Kelso J (2014) leeHom: adaptor trimming and merging for Illumina sequencing reads. Nucleic Acids Res 42(18):e141.

48. Guindon S, et al. (2010) New algorithms and methods to estimate maximum-likelihood phylogenies: assessing the performance of PhyML 3.0. Systematic biology 59(3):307–321.

49. Stamatakis A (2014) RAxML version 8: a tool for phylogenetic analysis and post-analysis of large phylogenies. Bioinformatics 30(9):1312–1313.

50. Drummond AJ & Rambaut A (2007) BEAST: Bayesian evolutionary analysis by sampling trees. BMC Evolutionary Biology 7:214.

51. Darriba D, Taboada GL, Doallo R, & Posada D (2012) jModelTest 2: more models, new heuristics and parallel computing. Nature methods 9(8):772.

52. Soares P, et al. (2009) Correcting for purifying selection: an improved human mitochondrial molecular clock. Am J Hum Genet 84(6):740–759.

53. Martfnez-Navarro B, Ros-Montoya S, Espigares MP, & Palmqvist P (2011) Presence of the Asian origin Bovini, Hemibos sp. aff. Hemibos gracilis and Bison sp., at the early Pleistocene site of Venta Micena (Orce, Spain). Quatern Int 243(1):54–60.

54. Qui Z, Deng T, & Wang B (2004) Early Pleistocene mammalian fauna from Longdan, Dongxian, Gansu, China. Paleontologia Sinica 191(C27):1–198.

